# Quantitative Analysis of the Optogenetic Excitability of CA1 Neurons

**DOI:** 10.1101/2023.06.02.543419

**Authors:** Ruben Schoeters, Thomas Tarnaud, Laila Weyn, Wout Joseph, Robrecht Raedt, Emmeric Tanghe

**Affiliations:** WAVES, Department of Information Technology (INTEC), Ghent University/IMEC, Technologypark 126, 9000 Ghent, Belgium; 4BRAIN, Department of Neurology, Institute for Neuroscience, Ghent University, Corneel Heymanslaan 10, 9000 Gent, Belgium

**Keywords:** Optogenetics, Cornu-Ammonis, Computational Modeling, NEURON, Tissue Activation, Sensitivity Analysis

## Abstract

Optogenetics has emerged as a promising technique for modulating neuronal activity and holds potential for the treatment of neurological disorders such as temporal lobe epilepsy (TLE). However, clinical translation still faces many challenges. This in-silico study aims to enhance the understanding of optogenetic excitability in CA1 cells and to identify strategies for improving stimulation protocols. Employing state-of-the-art computational models, the optogenetic excitability of four CA1 cells, two pyramidal and two interneurons, expressing ChR2(H134R) is investigated. The results demonstrate that confining the opsin to specific neuronal membrane compartments significantly improves excitability. An improvement is also achieved by focusing the light beam on the most excitable cell region. Moreover, the perpendicular orientation of the optical fiber relative to the somato-dendritic axis yields superior results. Inter-cell variability is observed, highlighting the importance of considering neuron degeneracy when designing optogenetic tools. Opsin confinement to the basal dendrites of the pyramidal cells renders the neuron the most excitability. A global sensitivity analysis identified opsin location and expression level as having the greatest impact on simulation outcomes. The error reduction of simulation outcome due to coupling of neuron modeling with light propagation is shown. The results promote spatial confinement and increased opsin expression levels as important improvement strategies. On the other hand, uncertainties in these parameters limit precise determination of the irradiance thresholds. This study provides valuable insights on optogenetic excitability of CA1 cells useful for the development of improved optogenetic stimulation protocols for, for instance, TLE treatment.

## I. Introduction

Optogenetics is a technique that can be used to manipulate cellular activity with light [1], [2]. Specific cell types are sensitized to light via the introduction of gene constructs that encode for optogenetic actuators [3]. Opsins are light-gated ion channels, pumps or receptors that when illuminated allow hyper-or depolarizing currents through the cell membrane [2]. The expression of these opsins is controlled by promoters expressed in the target cells [3]. Consequently, when applied to the brain, optogenetics provides the possibility to target specific neuronal populations which makes the technique highly promising to treat a variety of neurological disorders, such as epilepsy [4], [5].

Temporal lobe epilepsy (TLE) is one of the most common forms of epilepsy. 30% of patients with TLE cannot be helped with anti-epileptic drugs [6]. Based on the premise that temporal lobe seizures originate in the hippocampus, inhibition of the excitatory subpopulations is a possible strategy for TLE seizure suppression [7], [8]. Using the anion selective opsin halorhodopsin (NpHR) to inhibit hippocampal pyramidal cells has resulted in a decrease of seizure activity [9]–[11]. Excitation of the inhibitory hippocampal interneurons using cation conducting opsins, such as channelrhodopsin-2 (ChR2) and variants, has also yielded promising results [11]–[14]. Furthermore, optogenetics can be employed in studies on the initiation and propagation of seizures, and to investigate the role different neuronal populations play in the seizure dynamics [10], [15].

These studies are very promising but the development of clinical applications using optogenetics within the brain still faces some challenges [5], [6]. A first challenge is having a good understanding of the optogenetic effect, because aforementioned seizure-reduction strategies can have ictogenic effects as well [8], [12], [16]. Another challenge is bringing sufficient light into the brain. The optic frequencies have a poor penetration in brain tissue necessitating in-vivo implantation of the illumination source. In order to minimize structural damage due to intrusion, the light source’s dimensions are limited [17]. As a result only a small volume is illuminated with a single optical fiber, which could be insufficient for the human brain [8]. Furthermore, light absorption by the brain tissue causes heating. Therefore the light intensity should be limited in order to prevent brain damage [18]–[22].

To minimize stimulation power, one improvement could be the transition to red-shifted opsins [23], [24]. (Infra)red light gets less absorbed resulting in higher illumination volumes and less heating [19], [25]–[27]. Another solution is increasing the efficiency such that higher photocurrents are generated at lower irradiances. Improving single channel conductance or altering channel kinetics are possible strategies [24], [28], [29]. Another approach is enhancing membrane expression [3], [30], [31]. A final strategy is spatial confinement of the opsin to specific neuronal membrane compartments [3], [32]–[34].

Computational modeling can aid in developing a better understanding of the optogenetic response while avoiding in-vivo animal testing [36]. In 2006, a first mathematical description of the channel dynamics of ChR2 was published [37]. Since then, several studies have been published that use this and other, more accurate and efficient models in combination with neuron models to design in silico experiments [36], [38]–[43]. These experiments allow for easy parameter manipulation and exploration of the stimulation parameter space. Opsin expression can be spatially constrained, expression in different cell types can be tested, and interaction with the 3D light intensity profile can be evaluated. This optic field can be obtained via Monte Carlo based simulations [44] that simultaneously can be used to investigate the effect of tissue optical properties [18], [19], [22].

These optical field studies have shown that the light intensity spatial variation occurs on a neuronal scale. Studies combining optic field simulation with optogenetic response of neurons are scarce. Foutz et al. (2012) investigated this interaction with a multicompartment, neocortical layer V, pyramidal model [45]. Arlow et al. (2013) performed a study on a myelinated axon MRG model [46]. They identified a complex interplay of various simulation parameters on the activation threshold. On the other hand, Grossman et al. (2013) showed that opsin spatial confinement has an impact on action potential generation and propagation [47]. This non-uniform opsin expression has been touched upon by Foutz et al. (2012) as well. In this study, the optogenetic excitability of cornu ammonis (CA1) cells is investigated with the aim of gaining insights that could help guide optogenetic experiments concerning the suppression and initiation of seizures. The novelty is the use of morphologically reconstructed and data-driven biophysical models of CA1 pyramidal neurons and interneurons [35] that are extended to include ChR2(H134R) dynamics, and are subsequently subjected to a Monte Carlo simulated optic field. The effect of opsin expression level and spatial confinement on stimulation thresholds for multiple pulse durations is determined. Furthermore, the impact of various uncertain parameters, e.g., optical field properties, cell to optical fiber orientation and 3D structural cell morphology, is quantified. Based on these results, possible subcellular improvement strategies coupled with ideal optical fiber positioning are identified.

## II. Methods and Materials

The CA1 cell models used to test the optogenetic excitability are described below. Next, the light intensity fields determined via the Monte Carlo (M.C.) method and the opsin model are elaborated on. Finally, the methods and metrics used to analyze the optogenetic response are explained.

### A. Neuron models

Four models from Migliore et al. (2018) [35] are used in this study: two models of CA1 pyramidal cells and two CA1 interneurons located in the stratum pyramidale, i.e., a pravalbumin positive basket cell and a bistratified cell. All models have a different three-dimensional structural morphology, see figure 1 (A). The cell identification number of the pyramidal cells pyr_1_ and pyr_2_ is *mpg141208 B idA* and *mpg141209 A idA*, respectively. The bistratified cell (int_1_) has identification number *980513B* and the basket cell (int_2_) has number *060314AM2*. Although the pyramidal cells have a different structural morphology, they are classified under the same morphological type (m-type, [48], [49]). To avoid confusion, in this manuscript we will talk about difference in cell instead of morphology when comparing pyr_1_ vs. pyr_2_ (int_1_ vs. int_2_). Both pyramidal cells are continuous accommodating, while the electrical type or e-type of the interneurons is continuous non accommodating (Petilla convention [49], [50]). The models include ten active conductances: a sodium current, three types of potassium (K_DR_, K_M_, K_A_), three types of calcium (Ca_T_, Ca_L_, Ca_N_), two Ca^2+^-dependent potassium currents (K_Ca_, K_Cagk_) and a nonspecific current. Moreover, a simple calcium accumulation mechanism is included with a single exponential decay of 100 ms. The models were fitted to experimentally obtained voltage traces using a genetic algorithm. For more details of the models and the fitting procedure we refer to Migliore et al. (2018) [35]. The models themselves can be found on modelDB (https://senselab.med.yale.edu/modeldb/) under accession number 244688. The center of cell’s soma is at the origin. The somato-dendritic axis of the neurons is aligned to the z-axis. This somato-dendritic axis is the first principal component determined via a principal component analysis on the 3D-positions of all compartments except the axon. In case of the pyramidal cells, the apical trunk is always directed in the positive z-direction.

**Fig. 1.**
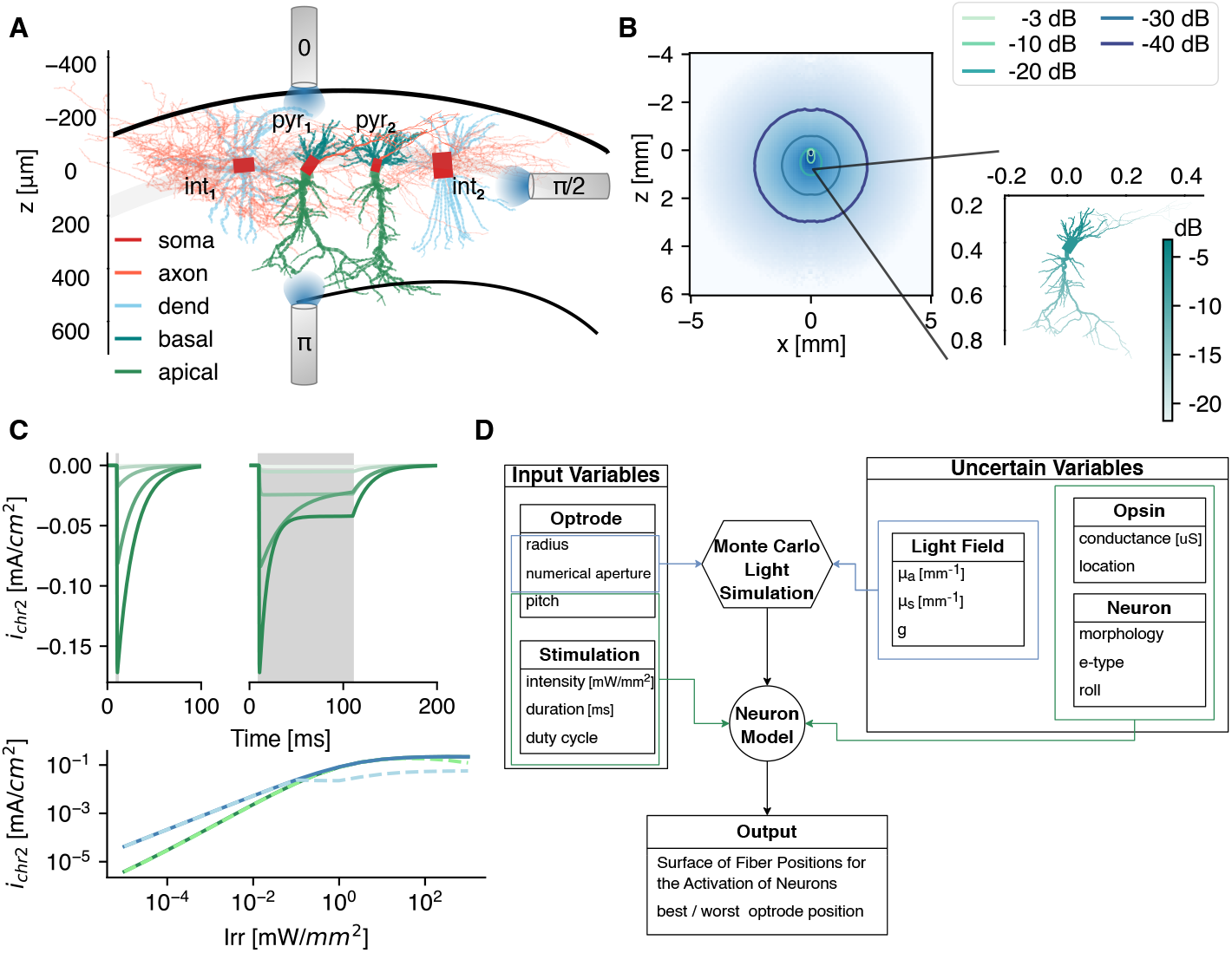
The simulation framework. **A** Graphical representation of the 3D reconstructed CA1 cell models used in this study: two pyramidal cells (pyr_1_ and pyr_2_), 1 bistratified (int_1_) and 1 basket cell (int_2_). The models are adopted from Migliore et al. (2018) [35]. This reference frame is applicable throughout the whole study, i.e. the soma is at z = 0 mm, and the somato-dendritic axis is parallel to the z-axis. As depicted, three optrode pitch setups are tested. The color represents the different opsin expression locations. **B** The light intensity profile in gray matter with *µ*_*a*_ = 0.42 mm^−1^, *µ*_*s*_ = 11.33 mm^−1^ and *g* = 0.88. The irradiance at the cell, with soma 400 *µ*m bellow the fiber, is shown in the inset. **C** The opsin current density at a voltage clamp of -70 mV for increasing irradiance (light to dark: 10^−3^→10 mW/mm^2^) of a 1 (left) and 100 ms (right) optical pulse, indicated in gray. (Bottom) The peak (solid) and steady state (dashed) current densities as function of irradiance for a 1 and 100 ms optical pulse in green and blue, respectively. **D** The simulation flowchart with input and uncertain parameters.

### B. Light Field in Gray Matter

To activate the opsins, the neurons are subjected to light. The light intensity field produced by an optical fiber (100 *µ*m radius, 0.39 numerical aperture (NA); after the optrode in the study of Acharaya et al. (2021) [18]) is determined via the Monte Carlo method. The used method is based on the direct photon flux recording strategy of Shin and Kwon (2016) [22]. The Monte Carlo simulations are solved in cylindrical coordinates for a homogeneous medium as in Stujenske et al. (2015) [19]. Because the hippocampus is predominantly a gray matter structure, the in-vivo optical properties of gray matter are selected. The absorption coefficient (*µ*_*a*_) at 470 nm is obtained by extrapolation of the data reported in Johansson 2010 [25] via a third order polynomial fit on the points between 480 and 550 nm. The scattering coefficient (*µ*_*s*_) and anisotropy factor (*g*) are obtained via interpolation from Yaroslavsky et al. (2002) [26]. This is often combined into the reduced scattering coefficient 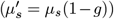. The refractive index (*n*) is 1.36. The simulations are run with 10^7^ photons and a radial (dr) and axial discretization (dz) of 5 *µ*m. The result of the Monte Carlo simulation with parameters given in table I is shown in figure 1 (B), with corresponding irradiance at the neuron level given in the inset. Throughout the study three pitch orientations with respect to the CA1 cells are tested as depicted in figure 1 (A).

**TABLE I.**
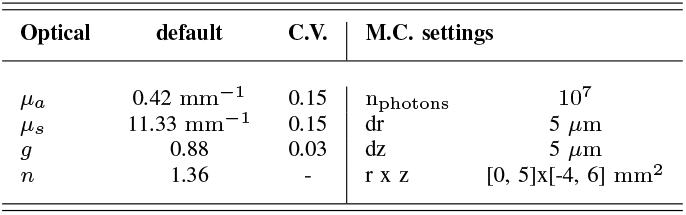
Parameters used in the Monte Carlo (M.C.) Simulations. Default values and coefficient of variation (C.V.).

### C. ChR2(H134R) Opsin

The selected opsin is a genetically engineered Channelrhodopsin-2 (ChR2) variant: ChR2(H134R). It has an enhanced steady-state photocurrent with a slower closing mechanism than the wildtype ChR2. Its peak operation is at a wavelength of 470 nm [28], [51]. The opsin’s kinetics are modeled with the double two-state opsin model, more specifically the RSRS-final reported in Schoeters et al. (2021), because of its improved computational efficiency compared to alternative four state Markov models [41]. It consists of two independent two-state pairs: *O* (opening and closing of the channel) and *R* (change in conductance due to dark-light adaptation). The transmembrane current density (mA/cm^2^) at a single section is determined as follows:

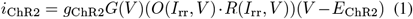

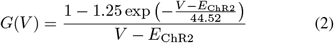

with *g*_ChR2_ the specific conductance (S/cm^2^), *G*(*V*) the rectification function given by equation (2), *V* the membrane potential (mV), *I*_rr_ the irradiance (mW/mm^2^) and *E*_ChR2_ the equilibrium potential (mV). The photocurrents for an optical pulse with duration of 1 and 100 ms with increasing irradiance under voltage clamp recording of -70 mV is shown in figure 1 (C).

The opsin is spatially confined to specific neuronal membrane compartments (in this manuscript further denoted as subcellular region). In case of the pyramidal cells, it is located in the soma, or distributed over the axon, basal dendrites (basal), apical dendrites (apic), all dendrites (dend = basal ∪ apic) or all sections (allsec = dend ∪ soma ∪ axon). In case of the interneurons, no distinction between apical and basal dendrites is made. The different regions are illustrated via the color code in figure 1 (A). In each simulation, *g*_ChR2_ is uniformly spread over the subcellular region of interest. Its value is calculated from a preset total maximum conductance:

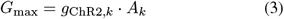

with *A*_*k*_ the total membrane surface area of a subcellular region *k*. The surfaces are summarized in Table II. *G*_max_ are 8 points spaced evenly on a log scale between 10^−1^ and 10^1.5^ *µ*S (for the uniform field 9 additional points are included between 10^−1^ and 10^2^).

**TABLE II.**
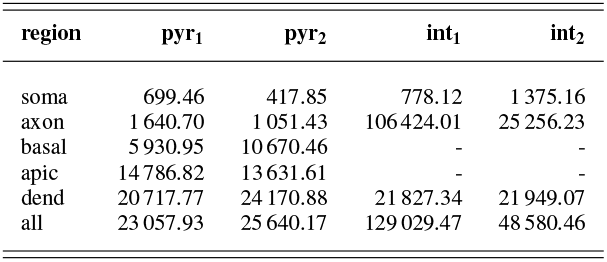
Total membrane surface area of subcellular regions in *µ*m^2^

Finally, the total temporal averaged current at the excitation threshold (TAC) [52] is calculated by:

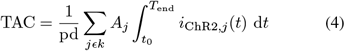

where pd is the pulse duration, *t*_0_ is the pulse onset time, *T*_end_ = max(500 ms, *t*_0_ + pd + 100 ms) (to ensure to include channel closure) and *j* is every compartmental segment in region *k*. The pd varied over 5 points logarithmic evenly spaced between 0 and 100 ms (9 pds *ϵ* [10^−1^, 10^3^] are used in the simulations with uniform light field).

### D. Analyses

The tests and metrics used to analyze the optogenetic response are described below.

#### 1) Surface of Fiber Positions for the Activation of Neurons

As metric to assess the optogenetic excitability, the surface of fiber positions for the activation of neurons (SoFPAN) is determined. This metric is similar to the more well-known volume of tissue activation (VTA) [53], [54]. In VTA, the frame of reference is the position of the stimulation device, while here, the frame of reference is the stratum pyramidale with the center of cell’s soma at origin. In other words, SoFPAN encompasses the positions at which the optical fiber can be located in order to activate the neuron of interest. This is chosen because of the layered structure of the hippocampus, constraining the cell bodies of the considered neurons to the stratum pyramidale. Moreover, the optical fiber is limited to the plane shown in figure 1 (A) to reduce the number of simulations.

The intensity threshold (*I*_th_) to elicit an action potential (V*>*-10 mV) recorded in the soma for a given pulse duration and fiber position is determined first. This threshold value is the intensity at the fiber surface (*I*_fiber_). It is obtained with a titration process using the bisection method for 7 iterations (i.e., *c*_*i*_ = (*a*_*i*_ + *b*_*i*_)/2, with *b*_0_/*a*_0_ = 10). 121 fiber positions are evaluated with z taking on eleven linearly spaced values in [-400, 700] *µ*m. For a pitch of *π*/2, x *ϵ* [-1000, 4000] *µ*m with Δx = 500 *µ*m. For the fiber pitch of 0 and *π*, x *ϵ* [0, 2500] *µ*m with Δx = 250 *µ*m. For a given pulse duration, the SoFPAN is then determined as the union of the spatial points for which the threshold is lower than the fiber intensity (true: *I*_th_ *< I*_fiber_) multiplied by the discretization surface (i.e., 110×250 or 110×500 *µ*m^2^ depending on the fiber pitch). A lower bound is determined by counting only the discretization surfaces enclosed by four true-points. In case of the upper bound, the true-false field is dilated first with a 3×3 mask before the enclosed-by-four-true-points regions are counted. For the pitches of 0 and *π*, the obtained SoFPAN is multiplied by two, such that the maximal SoFPAN for all pitches is equal to 5.5 mm^2^. The SoFPAN is calculated for nine logarithmic evenly spaced fiber intensities 0.1 and 1000 mW/mm^2^.

If the cell’s dimensions are assumed to be negligible with respect to the spatial variation of the intensity field, the irradiance is uniform over the whole neuron. The cell can thus be represented as a single point in space. The estimated SoFPAN under this assumption is denoted as SoFPAN_uniform_.

#### 2) Wilcoxon Signed-Rank Test

A single-sided, paired Wilcoxon signed-rank test is used to test whether the results from two populations are significantly different. For the uniform field 20 classes (cell and opsin location: 4×{all, axon, soma, dend} + 2×{basal, apic}) are mutually compared. In case of the M.C. field there are 42 classes (pitch, cell and opsin location: 3×[4×{all, axon, soma} + 2×{basal}]). A scoring system is used to identify the most excitable setup. Here, a 1, 0.1 and 0.01 is given if the SoFPAN is statistically greater with p-value below 0.001, 0.01 and 0.05, respectively. The maximum score is consequently 41.

#### 3) Regression

A two step fitting procedure is performed to analyze the effect of pulse duration and expression level on the intensity threshold as suggested by Williams and Entcheva [52]. First Lapicque’s formulation [55] is fit to the TAC. Second a linear regression is performed with independent variables the log_10_ of 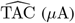 and *G*_max_ (*µ*S).

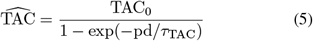

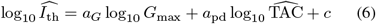

Here, TAC_0_, *τ*_TAC_, *a*_*G*_, *a*_pd_, and *c* are the regression parameters. 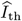 and 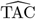 are the estimators of the threshold intensity and total averaged current, respectively.

#### 4) Optimal and Worst Fiber Position

The optimal and worst fiber positions for a given pulse duration are defined as the z-position where the depth of activation (amount of positions along the x-axis with *I*_th_ *< I*_fiber_) is the highest or the lowest, respectively. As a tie-breaker, the z-position is selected where the average of TAC along x is, respectively, the lowest or the highest.

#### 5) Elementary Effects

The inputs to the simulations can be divided into two categories (figure 1 (D)). On the one hand, there are the parameters that are known (e.g, optrode radius, NA and pitch) or controlled by the user (e.g., pd and *I*_fiber_) in an optogenetic experiment. On the other hand, there are variables that cannot be controlled and contain some uncertainty, e.g., the tissue’s optical properties 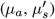, the neuron’s structural morphology (cell: pyr/int_1_ vs. pyr/int_2_) and orientation (roll, yaw), and the opsin expression (level (*G*_max_) and location). Because of the expected non-linear interactions and high computational time (*±*12 hours for a 121 position sweep with 5 pulse durations), the elementary effects method is adopted for global sensitivity analysis [56].

Six influential factors are investigated, i.e., 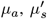, *G*_max_, cell, opsin location and roll. The measure used (*µ*^∗^) is the mean of the absolute value of *r* = 16 elementary effects. Also, the standard deviation of the elementary effects (σ) is calculated to track interactions and non-linear effects. The *r* repetitions are sampled with the radial-based design using Sobol’s numbers [57]. 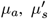 and *G*_max_ are normally distributed with mean and coefficient of variation (C.V.) given in table I. The mean and C.V. of *G*_max_ are 1 *µ*S and 0.15. The roll is uniformly distributed between [-*π, π* [radians. Cell and location are discretely uniformly distributed, with {pyr_1_, pyr_2_} ({int_1_, int_2_}) and {all, axon, soma, basal} ({all, axon, soma}) classes, respectively, for pyramidal (inter-) neurons.

### E. Software

Simulations were done with NEURON 8.0.0 [58] and Python 3.9.12. on the HPC system with AMD Epyc 7552 processing units, provided by the Flemish Supercomputer Center.

## III. Results

The results of this study can be subdivided into three sections. First, there is the optogenetic response under a uniform field. This means that all cell sections receive the same irradiance. This facilitates the analysis of the importance of subcellular regions, pulse durations and expression levels. Next, the Monte Carlo simulated light field is included. This allows us to indicate the effect of light propagation on optogenetic excitability and provides information on optimal or worst fiber positioning. Finally, an elementary effects study is performed to identify the most sensitive parameters.

### A. The Optogenetic Excitability in a Uniform Light Field

Figure 2 (A) depicts the intensity threshold (*I*_th_) and temporal averaged current (TAC) curves as function of pulse duration (pd) of several subcellular and cell combinations. Both decrease with increasing pd until a saturation value is reached. For the same subcellular region (i.e., allsec), the threshold curve is shifted down in case of pyr_1_ versus int_2_ (blue solid vs. blue dashed-dotted line). Staying within the same cell, it can be seen that changing the subcellular region (all → basal or soma) can result in a downward shift, as well. The same effects can be seen on the TAC, but less pronounced. Increasing the expression level (*G*_max_) from 1 to 10 *µ*S (blue solid line with circle markers) results in an expected decrease of the threshold. On the other hand, the TAC remains the same. This is highlighted in figure 2 (B) where the TAC is constant for all pds for a *G*_max_ *>* 1 *µ*S in the allsec-pyr_1_ setup. A linear relationship for *I*_th_ can be observed for *G*_max_ ≥ 0.518 *µ*S. No threshold could be found for *G*_max_ *<* 0.215 *µ*S. This is because the photocurrent saturates at high intensities (see figure 1 (C)) and therefore cannot compensate for the low *G*_max_. At high pulse durations a decrease in TAC is already observed at higher *G*_max_ (*<* 1 *µ*S). For the corresponding *I*_th_, the photocurrent is biphasic with a peak and steady-state value. At these pulse durations (100 and 1000 ms) the action potential (AP) occurs during the inactivation from peak to steady state. The occurrence of the peak is much more efficient in eliciting an AP. A large part of the total photocurrent occurs before the AP while this diminishes at higher *G*_max_ with monophasic photocurrents. Additionally, during the AP, the current drops to zero due to channel shunting. Consequently, the net effect of this drop on the TAC is larger here than in case of no biphasic photocurrents (at low irradiances) or lower pulse durations, where the AP is elicited during the deactivation phase of the photocurrent.

**Fig. 2.**
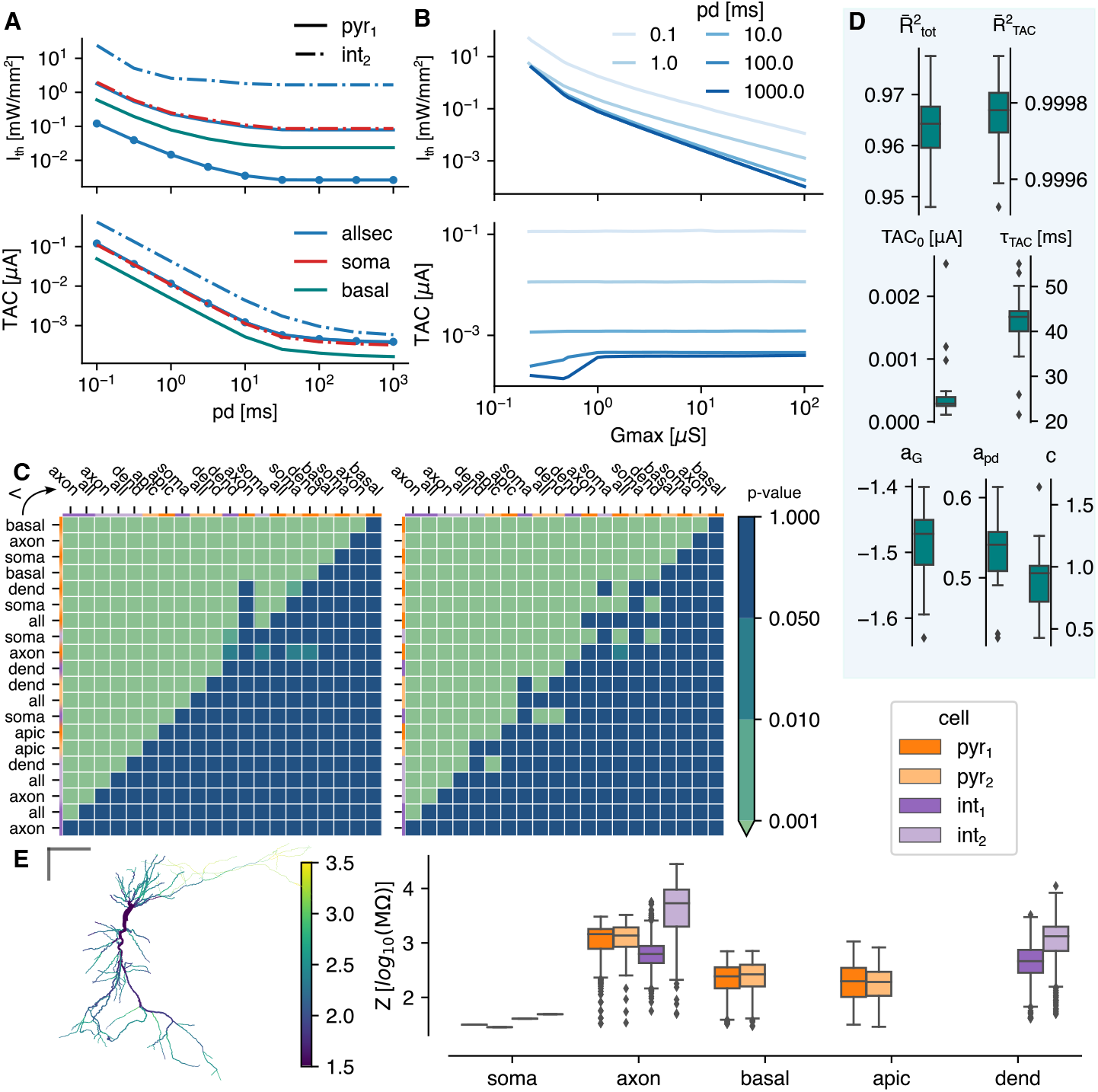
The optogenetic response under a uniform light field. **A** The intensity threshold (*I*_th_) and corresponding total temporal averaged current (TAC, top and bottom, resp.) as function of pulse duration. All lines shown are for a *G*_max_ = 1 *µ*S except the circle markers (*G*_max_ = 10 *µ*S). **B** The intensity threshold and corresponding total temporal averaged current (top and bottom, resp.) as function of *G*_max_ for allsec-pyr_1_. **C** The p-value of the mutually, paired Wilcoxon signed-rank tests. Single-sided test if class at the row is lower than class in the column for the *I*_th_ (left) and TAC (right). The color code at the top and left indicate the cell model. **D** The results of the two step regression. The goodness of fit is indicated with the adjusted R-squared measure: 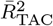 for the Lapicque fit to the TAC and 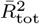 for the linear regression fit to the threshold. Box plots are shown for 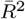 -values and regression parameters, calculated for the different cell-types and opsin locations. **E** Summary of the input impedance at 0 Hz, (left) the local impedance of each section in pyr_1_, (right) the summarized impedances of the four tested cells. The gray bars are 100 *µm*

Mutually paired Wilcoxon singed-rank tests are performed to identify the most excitable subcellular region. The p-values of a single-sided test are shown in figure 2 (C) of the threshold intensity and TAC on the left and right, respectively. Each compared population consists of 153 points (9 pds and 17 *G*_max_ values). The basal-pyr_1_ is the most excitable with a *I*_th_ significantly (p*<*0.001) lower than any other location-cell combination. The same is true for the TAC. The pyramidal cells are more excitable than the interneurons with pyr_1_ the most excitable (p*<*0.001). Aside from the axon in pyr_2_, the order of the excitability scores is the same in both cells (see section II-D2). The apical dendrites is the least excitable subcellular region of the pyramidal cells (p*<*0.001). On average a lower *I*_th_ is required for the basket cell than the bistratified cell (p*<*0.001). Overall a significantly lower threshold is accompanied with a significantly lower TAC. Some exceptions exist where the TAC is significantly greater (e.g., all-pyr_1_ versus soma-int_2_). The median of the relative change in *I*_th_ of the location-cell combinations as ranked in figure 2 (C) is shown in figure S.1 (A). The average of the medians is - accompanied with a significantly lower TAC. Some exceptions exist where the TAC is significantly greater (e.g., all-pyr_1_ versus soma-int_2_). The median of the relative change in *I*_th_ of the location-cell combinations as ranked in figure 2 (C) is shown in figure S.1 (A). The average of the medians is - 22.76%. Therefore, the *I*_th_ drops every time on average with 22.76% percent when moving from the least (axon int_1_) to the most (basal pyr_1_) excitable location-cell combination. Optimal subcellular expression in a cell can result in *I*_th_ drops of *>*75%. Compared to no subcellular specificity (opsin over the whole cell), *I*_th_ reductions of *>*60% can be achieved in the pyramidal cells, increasing towards 83% and 92% for the basket and bistratified cell, respectively (cf. table III).

**TABLE III.** Influence of subcellular opsin expression. Median relative change between best and worst subcellular location, and best and no subcellular specificity (all) of all pd And *G*_max_ Combinations. excitability under a uniform light field (*i*_th_) and with light propagation (SOFPAN).

| cell | $I_{th}$ | | | SoFPAN | | |
| --- | --- | --- | --- | --- | --- | --- |
|  | best | worst | % | best | worst | % |
| pyr1 | basal | apic | -81.23 | basal $\pi/2$ | axon 0 | 223.08 |
| | basal | all | -67.63 | basal $\pi/2$ | all $\pi/2$ | 33.33 |
| pyr2 | axon | apic | -76.03 | axon $\pi$ | all 0 | 152.78 |
| | axon | all | -64.36 | axon $\pi$ | all $\pi$ | 67.42 |
| int1 | alldend | axon | -99.89 | soma $\pi/2$ | axon 0 | 947.62 |
| | alldend | all | -91.58 | soma $\pi/2$ | all $\pi/2$ | 100.00 |
| int2 | soma | axon | -87.28 | soma $\pi/2$ | axon 0 | 233.33 |
| | soma | all | -82.84 | soma $\pi/2$ | all $\pi/2$ | 87.47 |

The two step regression is performed on each location-cell combination separately. The combined results are shown in figure 2 (D). The variability on TAC is captured by Lapicque’s formulation resulting in a median adjusted coefficient of determination 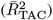 value of 0.99978. The median rheobase (TAC_0_) is 3.17 nA and the median time constant (*τ*_TAC_) is 42.95 ms. The variability of *I*_th_ is well explained by the two step regression model 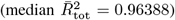. The median parameter values of *G*_max_ and pd (*a*_*G*_ and *a*_pd_) are -1.47 and 0.53, respectively. Even though the latter is positive, *I*_th_ decreases with pd due to the Lapicque’s formulation. Based on these values, *G*_max_ has thus a stronger impact on *I*_th_ than the pd.

The input impedance of the cell’s sections are shown in figure 2 (E). The impedance is a proportionality factor on the input current resulting in a given voltage change. Therefore, the voltage change is higher for a higher input impedance given a constant input current. The axons have on average the highest impedance but also the highest spread with many outliers. At the left, it can be seen that the impedance increases from axon hillock to the synapses. The soma has a low impedance compared to the median of the other subcellular regions.

### B. The Optogenetic Response in the Monte Carlo Light Field

In this section, the cells are subjected to an intensity field produced by a 100 *µ*m fiber. The 3D-profile is obtained via the Monte Carlo method in homogeneous gray matter tissue with the default optical parameters as summarized in table I. Unlike in the previous section, the irradiance at the different neuron sections will depend now on their relative position with respect to the fiber and its orientation. Consequently, *I*_th_ now depends on the position in the reference frame as indicated in figure 1 (A). An example is shown in figure 3 (A), with the *I*_th_ map on the left and corresponding TAC on the right, for an allsec-pyr_1_ with *G*_max_ = 1.18 *µ*S, pd = 10 ms and a fiber pitch of *π*/2. A hotspot is observed around [*x* = 0.43 mm, *z* = −0.036 mm], where the threshold is the lowest. Even when the neuron lies behind the fiber, i.e., *x <* 0 mm, it is still possible to excite the cell but at higher *I*_th_. Two islands can be observed for the TAC with a factor two difference between minimum and maximum. The farther away from the cell the lower the variation in TAC.

**Fig. 3.**
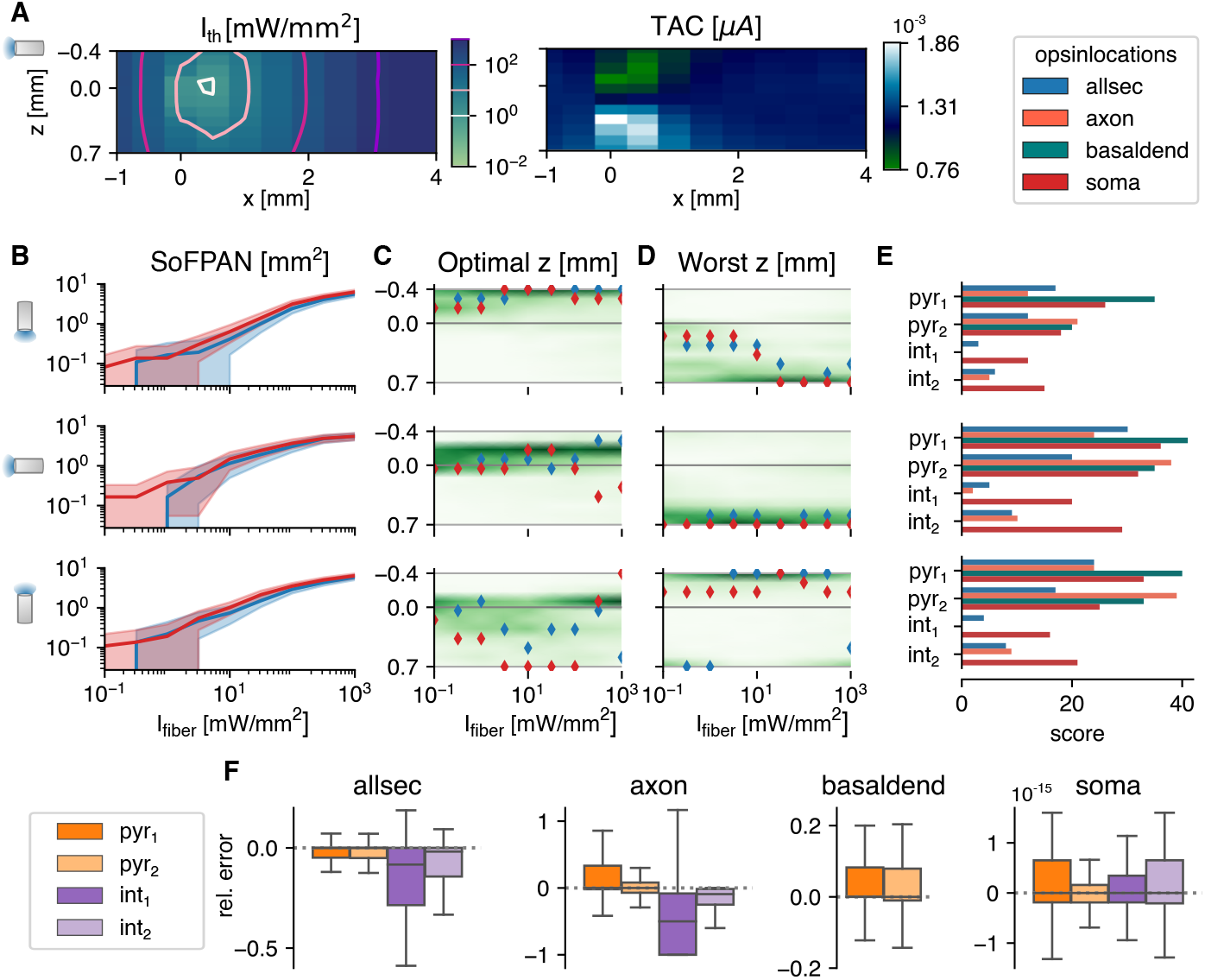
The optogenetic response in a homogeneous Monte Carlo simulated light field. **A** The threshold intensities (left) of the pyr_1_ cell, with opsin distributed over all sections, and corresponding total temporal averaged current (right). In the colormaps, the position of the optical fiber is varied with respect to the soma, which is fixed at (*x, z*) = (0, 0). **B, C and D** The Surface of Fiber Positions for the Activation of Neurons, the optimal and worst z-position for neuron activation, respectively, as function of fiber intensity. The shaded area in **B** is enclosed by the upper and lower bound. The lines are the result of the pyr_1_ cell with opsin location shown in legend at the right top corner. *G*_max_ = 1.179 *µ*S and pd = 10 ms in **A-D**. In **C and D** the scatter plots overlay the density, averaged over all cell, location, *G*_max_ and pd combinations. **E** Excitability score based on paired Wilcoxon signed-rank tests. Each population consists of 360 (5 pd × 8 *G*_max_ × 9 *I*_fiber_ values) combinations. The pitch of the fiber is illustrated by the orientation of the fiber icon on the left and valid for the whole row. **F** Relative error of SoFPAN, if the light intensity is considered uniform over the neuron: rel. error = (SoFPAN_uniform_ - SoFPAN_M.C._)/SoFPAN_M.C._, the outliers are not shown. The errors are calculated for all the *G*_max_, pd, pitch and *I*_fiber_ combinations.

Based on these threshold maps, the surface of fiber positions for the activation of neurons (SoFPAN) can be estimated. This is shown in figure 3 (B) for the three investigated fiber pitches. The SoFPAN is a measure of the excitability of the subcellular optogenetic configuration in the light field. The uncertainty caused by the discretization on the SoFPAN is indicated by the shaded area (see section II-D1). There is a larger uncertainty at lower intensities due to the rough simulation grid. The soma appears to be more excitable than the opsin in all sections (red vs. blue) with already a non-zero SoFPAN for a *I*_fiber_ of 0.1 mW/mm^2^. This is true for all fiber pitches. The SoFPAN saturates due to the finite simulation domain.

An optimal and worst z-position for each fiber pitch can be determined, as well. The result for the two cases as above are shown in figure 3 (C, D). It can be seen that the positions vary with increasing *I*_fiber_. At low intensities, the optimal position is near the subcellular region (cf. soma, red) or the most excitable region (cf. allsec, blue), which is the basal dendrites. At higher intensities, it is better to illuminate the whole simulation domain, explaining the optimal positions at - 0.4 mm and 0.7 mm for the 0 and *π* pitches, respectively. Once the whole simulation field is excited, the optimal position is completely determined by the average TAC along the x-axis. This can be seen in the sudden changes at the highest *I*_fiber_ (*>* 100 mW/mm^2^). In green the density distribution of all 560 combinations (pd × *G*_max_ × loc × cell) is displayed. The optimal position is more concentrated with a 0 or *π*/2 pitch. On the other hand, the worst position is more concentrated with the *π*/2 and *π* pitches. The optimal and worst fiber positions are most distant with the *π*/2 pitch position.

The excitability of the subcellular region is summarized in figure 3 (E). The scores are based on the p-values of the single-sided, paired Wilcoxon signed-rank test, as described in II-D2. At each pitch individually, the subcellular excitability order remains, generally, the same as under the uniform field (cf. figure S.2). Only the axons appear to be susceptible to the fiber pitch position. For instance, the axon-pyr_1_ combination has an increased (decreased) excitability under the *π* (0) fiber pitch. Also at pitch 0, the axon of pyr_2_ has a lowered excitability with a SoFPAN significantly lower (p*<*0.001) than soma-pyr_1_. On the other hand, axon-int_2_ has a higher SoFPAN than all-int_2_ under the *π*/2 and *π* pitches. Basal-pyr_1_ is under all pitches the most excitable combination, with a score of 41 at pitch *π*/2. Overall, pitch *π*/2 is significantly more excitable than *π*, which is in turn more excitable than 0 (p*<*0.001). The axon-pyr_2_, however, has a significantly higher SoFPAN (p*<*0.001) under pitch *π* than *π*/2. The median of the relative change in SoFPAN of the location-cell-pitch combinations according to this ranking, i.e., lowest to highest score, is shown in figure S.1 (C). The average of medians is now 10.45%. Therefore, the SoFPAN increases every time on average with 10.45% percent when moving from least (axon int_1_, pitch = 0) to most (basal pyr_1_, pitch = *π*/2) excitable location-cell-pitch combination. The same analysis is performed when restricted to a single pitch. The average of medians of the relative changes in SoFPAN increase towards 56.86, 25.89, 59.18% for a pitch = 0, *π*/2 and *π*, respectively. Matching the ideal fiber location to the subcellular expression of a single cell can result in doubling of the SoFPAN compared to the worst combination. Even a tenfold increase is observed in case of the bistratified cell. Compared to subcellular unspecificity (all), SoFPAN increases of 33-100% can be obtained by specifying subcellular expression (cf. table III).

The SoFPAN for a light field that is considered to be uniform over the whole cell (SoFPAN_uniform_, see bottom section II-D1) is calculated, as well. The relative error compared to the SoFPAN under a Monte Carlo field, i.e., (SoFPAN_uniform_-SoFPAN_M.C._)/SoFPAN_M.C._, is shown in figure 3 (F). The errors is calculated for all the *G*_max_, pd, pitch and *I*_fiber_ combinations. The error of the soma is negligible. The outliers are not shown. Of these, only 0.85% produce a relative error above 5%. These results validate the method as the soma should not depend on the M.C. field as its section is only one point in the 3D-space. The SoFPAN of basal dendrites gets overestimated, with 18.17% having a relative error above 20%. In case of the pyramidal cells and with the opsin distributed over the whole cell, the SoFPAN is predominantly underestimated and the interquartile range (IQR) of the relative error is 60% of the IQR for the basal dendrites. The estimation for the axon of the int_1_ cell is the worst with a median relative error of -50%. At least in one test case of each subcellular region the error is either -100% or infinity (see table IV).

**TABLE IV.** Summary relative error Sofpan monte carlo versus Sofpan uniform (all cells pooled together). +20% indicates amount of simulations with a relative error above 20%

|  | outliers [%] | +20% [%] | median | max | min | IQR |
| --- | --- | --- | --- | --- | --- | --- |
| allsec | 17.42 | 23.96 | -7.8e-16 | ∞ | -0.59 | 0.12 |
| axon | 18.60 | 34.60 | 0 | ∞ | -1 | 0.20 |
| basal | 20.44 | 18.17 | 1.3e-15 | ∞ | -0.53 | 0.09 |
| soma | 24.57 | 0.17 | 0 | ∞ | -1 | 5.4e-16 |

### C. Parameter Uncertainties

There is uncertainty on various parameters used in this study. The optical parameters depend on multiple factors, e.g., tissue and wavelength. The absorption coefficient is extrapolated which introduces an uncertainty, as well. Moreover, the values of gray matter are used while different gradations exist. The effect of a change in optical parameters on the light field in homogeneous tissue, is illustrated in figure 4 (A). On the left, the field as used in the section above is depicted. The effect of an increased absorption and decreased reduced scattering coefficient is shown in the middle. The result is a more conical field with higher degradation. A more round field is obtained when the reduced scattering coefficient is higher (cf. right). Also at the cell level, there are multiple sources of uncertainty. In experimental setting, the opsin expression *G*_max_ will not be exactly known. Moreover, the subcellular location will probably not be discrete as used in these simulations. Finally, the morphology of the tested cells and its orientation (cf. roll) are fixed. To address the impact of these uncertainties on the output, a global sensitivity analysis is performed. The used approach is a screening method: the Elementary Effects test.

**Fig. 4.**
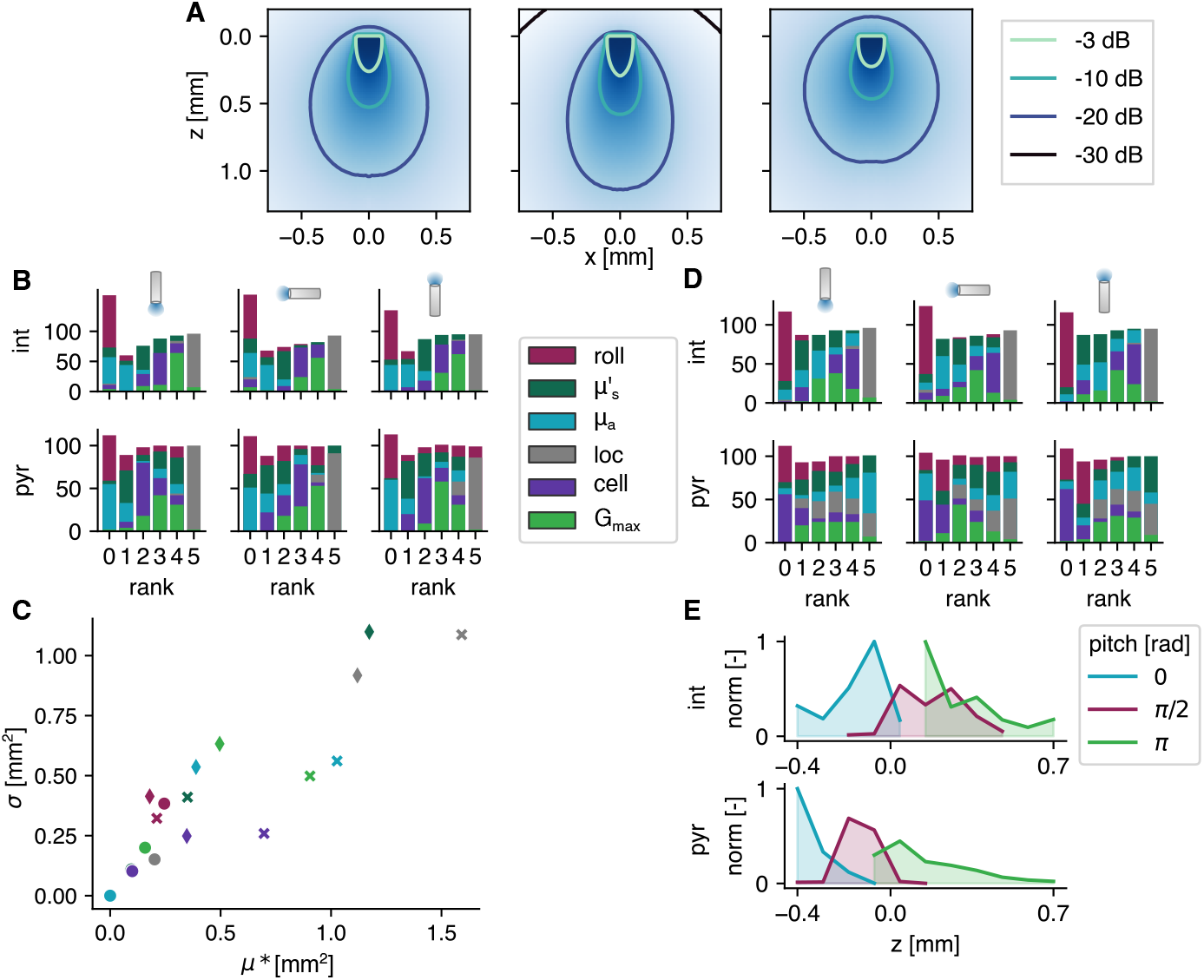
Results of the elementary effects study. **A** Intensity fields with different optical parameters; from left to right *µ*_*a*_ = 0.42, 0.52, 0.35 mm^−1^ and 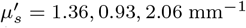. **B and D** The influential parameters on SoFPAN and optimal fiber position, respectively, ranked for 5 pds and 9 fiber intensities (in %). The fiber pitch and cell type are shown on top and left, respectively. **C** The two measures of the elementary effects of three pyramidal cell setups, i.e., pitch = *π, π*/2 and 0, and *I*_fiber_ = 1, 1000 and 1000 mW/mm^2^ indicated by circle, diamond and cross, respectively, for a pd = 10 ms. **E** Normalized optimal z-position over all results with allsec subcellular location of the elementary effects study.

The influence of these six parameters, i.e., cell roll, 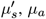, opsin subcellular location (loc), cell model (cell) and *G*_max_, on the SoFPAN for the three fiber pitches and two cell types (pyr and int) is investigated. For each fiber pitch and cell type, the elementary effects test (EET) is repeated for 5 pd and 9 *I*_fiber_ combinations (these 45 combinations correspond with 100% in figure figure 4 (B, D)). The rank according to the *µ*^∗^-measure is summarized in figure 4 (B). For certain pd and *I*_fiber_ no differentiation could be made based on the *µ*^∗^-measure, explaining the bar height *>*100% at rank 0. The subcellular location has most frequently the highest impact on the excitability for all six test cases. This is followed by *G*_max_ in second place (rank 4). The roll and *µ*_*a*_ have on average the lowest impact. However, in case of the pyramidal cells and *π* fiber pitch, the roll has occupied the highest rank for some (pd, *I*_fiber_) combinations. On the other hand, 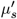 is more important when the fiber pitch is 0 for the pyramidal cells. The *µ*^∗^ and σ measures of three cases of the pyramidal cells with pd of 10 ms are shown in figure 4 (C). The circles indicate the setup where the roll has the highest rank, i.e., pyramidal cell with *π* pitch and 1 mW/mm^2^. It can be seen that even though it has the highest rank, its measures are in the same range as the other two cases. The diamonds represent the EET of a *π*/2 pitch at *I*_fiber_ = 1000 mW/mm^2^. Here, the reduced scattering coefficient has the highest impact. The non-linear and interaction effects are higher in this case, reflected by the higher σ. For the 0-pitch with *I*_fiber_ = 1000 mW/mm^2^ (cross), the location is ranked highest with a clear difference in *µ*^∗^. The effect of cell appears to be more linear than the other parameters indicated by its shift towards lower σ values.

The same analysis is performed on the optimal position. The rank is summarized in figure 4 (D). For the interneurons, the subcellular location stays dominant. There is, however, a clear shift towards the optical parameters for the pyramidal cells. Here, the reduced scattering and absorption coefficient are most often ranked highest for the *π* and 0 pitch, respectively. Also, in case of the *π*/2 pitch, the absorption coefficient is more important for the optimal position than it is for the SoFPAN. The cell and roll appear to have the lowest effect. While, on the other hand, the cell is important in case of the interneurons. The optimal position for both cell types normalized over all simulations with allsec subcellular location of the EET study, is shown in figure 4 (E). These exclude the preset subcellular selectivity. For the interneurons this is more smeared out and a focus towards the stratum pyramidale is observed. On the other hand, for the pyramidal cells there is a clear preference for a position such that the majority of the light reaches the stratum oriens region. At pitch *π* this is more smeared out due to the possibility to retract (z more positive) the fiber at higher intensities to illuminate a bigger region in the stratum oriens. This is limited for the 0 pitch explaining the high peak at −0.4 mm.

## IV. Discussion

In this study we focused on the optogenetic excitability of CA1 cells. We attempted to not only gain more insight into the effect of the various stimulation and uncertain parameters but also to identify strategies for increased optogenetic efficiency. These insights are of interest for the development of better stimulation protocols that can be used as treatment for TLE. A broad view is adopted where both the excitability of pyramidal and interneurons is investigated. Even though excitation of inhibitory neurons is one of the two main investigated strategies as treatment of TLE, insights in the excitability of pyramidal neurons can be of interest as well. Like with electrical deep brain stimulation, the latter could be used as counter-irritation [4] with various modes of action [59] that can be tested. Moreover, stimulation of both types could be beneficial for restoring the excitation-inhibition balance [16].

### A. Excitability of Spatially Confined Opsin Expression

The results show that the optogenetic excitability of CA1 cells depends on various parameters. The irradiance threshold ranges over multiple order of magnitudes. As expected, an increase in expression level (*G*_max_) or pulse duration (pd) results in a decrease of the intensity threshold (*I*_th_). There is also a clear dependence on the subcellular region of opsin expression and variance among different cells. There is no single explanation for the relative excitability of the considered subcellular opsin locations, due to the complex interplay of many non-linear relationships. By comparing the membrane areas and impedances (cf. table II and figure 2 (E)) of the subcellular regions some observations can be made. For a fixed *G*_max_, the specific channel conductance (*g*_ChR2_) is locally higher for regions with a lower total surface area. Thus, for the same *I*_fiber_, there will locally be a higher depolarizing current to elicit an action potential (AP). This combined with the fact that the AP is measured in the soma (therefore does not have to travel through the cell), explains why the soma-confinement is highly ranked in each cell. The observation concerning the locally raised channel conductance also holds when comparing basal dendrites with apical, all dendrites and all sections. An argument for why confinement to the basal dendrites is more excitable than the soma could be found by comparing their input impedances. For a fixed depolarizing current, a higher impedance results in a larger membrane depolarization. Because the impedance of the basal dendrites is significantly higher than that of the soma (log-scale in figure 2 (E)) it will facilitate AP initiation. However, this contradicts the rank of axon-pyr_1_. Finally, there is the channel distribution inside the cell itself. The ratio of depolarizing (e.g., Na^+^ and Ca^2+^) and hyperpolarizing (e.g., K^+^) channels defines the membrane threshold for AP initiation. This ratio is double in the axon of pyr_2_ compared to pyr_1_, motivating the low rank of axon-pyr_1_. These observations are in agreement with the findings of Foutz et al. (2012) [45].

Confinement to the basal dendrites of pyr_1_ is the most excitable, while the highest *I*_th_ is required for the axon of int_1_ (cf. figure 2 (C)). Similar ranking is observed in the SoFPAN calculations (cf. figure 3 (E)). To identify the effect of endogenous channel distribution, the channel distributions of pyr_1_ were imposed on pyr_2_ (pyr_3_) and vice versa (pyr_4_). The mean relative difference of all pd and *G*_max_ combinations is shown in table V. Compared to pyr_1_, the *I*_th_ drops with ∼30% for all subcellular regions except axon and basal dendrites. Contrarily, the excitability drops (higher *I*_th_) when the channel distribution of pyr_2_ is imposed on pyr_1_, except for the basal dendrites. In both cases, the *I*_th_ of the axon subregion almost doubles. It is clear that the interaction of structural morphology and channel distribution has an impact on optogenetic excitability. Neuron degeneracy, i.e., the ability to perform the same functioning whilst being structurally different or having different ion channel distributions [35], is thus something that should be taken into account in determining irradiance thresholds. Still, the basal dendrites region is also the most excitable in pyr_3_ and pyr_4_. Combined with rank 1 and 2 for pyr_1_ and pyr_2_, respectively, it can be concluded that this is the most effective subcellular target region for opsin expression in CA1 pyramidal cells.

**TABLE V.** Mean relative difference in *i*_th_ by changing channel distribution (relative with respect to original).

|  | all | alldend | apic | axon | basal | soma |
| --- | --- | --- | --- | --- | --- | --- |
| pyr<sub>3</sub> - pyr<sub>1</sub> | -0.32 | -0.31 | -0.28 | 0.83 | 0.14 | -0.31 |
| pyr<sub>4</sub> - pyr<sub>2</sub> | 0.29 | 0.23 | 0.47 | 0.92 | -0.28 | 0.30 |

This spatial dependence was also observed in the study of Grossman et al. (2013) [47]. With the specific conductance (*g*_ChR2_) as metric, they determined whole cell illumination to be most efficient, i.e., uniform opsin distribution over whole cell with a uniform light field compared to soma, axon initial segment or apical dendrite confinement. They determined that for a 20 ms pulse and irradiance of 1 mW/mm^2^ the required *g*_ChR2_ when in all sections is only 6% of that when restricted to the soma. On the other hand, *G*_max_ was 60% higher. These values are in agreement with our results where the ratio of the specific conductance of all sections to soma targeted expression is 2-4% under the same conditions in the pyramidal cells. However, in our study we argue that ranking should be based on *G*_max_, i.e., where the number of opsin complexes is fixed. This translates towards an equal comparison of total elicited photocurrent, while, on the other hand, for a fixed *g*_ChR2_ the total photocurrent is scaled by the surface area. As a result, the confinement to the soma is classified here to be more excitable.

After correction for the difference in rectification function (i.e., *G*(*V* = −68.83 mV) = 1 in [40] vs. 0.07 in this study), the absolute values of *G*_max_ were slightly lower but in the same order as reported in [47]. For a *G*_max_ of 1 *µ*S (= 0.07 *µ*S after correction) and with a single channel conductance of 40-100 fS this translates towards expression of 0.71-1.77 10^6^ opsin complexes. Spread over the whole cell this is *±*50 channels/*µ*m^2^ but confined only to the soma this rises towards *>*1000 channels/*µ*m^2^. This value is higher than the estimated 130 channels/*µ*m^2^ based on bacteriorhodopsin expression [45], [60] but lower than the indirectly estimated 4.4 10^4^ channels/*µ*m^2^ by Arlow et al. (2013) [46]. With our tested *G*_max_ values up to 100 *µ*S especially when restricted to the soma, this could pose cellular toxicity problems, if these channel numbers would be achieved [3], [24], [32]. This can be avoided if single channel conductance is increased.

In-vivo this highly specific and discrete separation is not possible. Still, by merging the opsin with signaling and targeting constructs, localized enhancement can be obtained. For instance, the addition of the soma-targeting motif of the soma- and proximal dendrite-localized voltage-gated potassium-channel Kv2.1 improves somato-dendritic expression [3], [33]. Our results encourage this approach and advances in this research direction. A reduction of *I*_th_ with more than 64% can be achieved via subcellular specificity. This is tempered towards, but still significant, increases in SoFPAN of 33-100%, when light propagation is included. Consequently, if made possible, spatial confinement of opsins to specific membrane compartments could significantly increase optogenetic efficiency. On the other hand, more than 76% reduction of *I*_th_ between optimal and worst subcellular regions is possible. This is also reflected in SoFPAN, where an ideal subcellular-pitch combination can result in a 1.5-10 fold increase compared to the worst combination (cf. table III). A good knowledge of the optogenetic interaction at the subcellular level is therefore required in order to achieve the optimal configuration.

### B. Optimal Fiber Position

Due to the finite size and discrete nature of the test grid (121 points), the SoFPAN, and optimal and worst positions could not be unambiguously determined. An upper and lower bound of the SoFPAN is defined (cf. sec. II-D1), indicating a larger uncertainty at lower intensities. In case of the optimal fiber position, a tie-breaker based on the TAC is introduced. From a homeostatic point of view, it is ideal that the required result is achieved via the lowest perturbation of normal functioning. High/long transmembrane currents could lead to ion concentration imbalance. Especially when using ChR2(H134R), which has a high H^+^ permeability, this can result in neuron acidification, which in turn can result in decreased neuron functioning or unexpected behavior [39], [61], [62]. Therefore, the positions that generate the lowest TAC are preferred. The results in figure 3 (C) show that the optimal position is at the depth of the region of subcellular expression or with a focus on the most excitable region (in case of opsin distribution over the whole or majority of the cell). Overall, the rank of excitability between uniform and M.C is unchanged. field stimulation. Subcellular excitability appears to be dominant over spatial distribution. This spatial preference was also observed in Foutz et al. (2012). In their L5 pyramidal model, they found the apical tuft and soma to be most excitable [45].

For the *π*/2 pitch, the optimal and worst position are the most stationary for increasing intensities. For the other pitches (0 and *π*), either the optimal or worst position is more smeared out while they are located more closely to each other for low intensities. Consequently, there is a higher risk for sub-optimal fiber positioning. Combined with the highest excitability according to the SoFPAN (cf. figure 3 (E)), it can be conclude that *π*/2 is the better fiber position.

### C. Contribution of Optical Field Simulation

This study combined simulations of light propagation and neuronal modeling. Light propagation is simulated using the Monte Carlo method for a uniform medium. The hippocampus is a predominantly gray matter structure. However, there is uncertainty on the exact values of the optical parameters (amplified by inter- and extrapolation). The effect of the uncertainty of these parameters is tested using the elementary effects method. The influence of 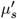 on the excitability is ranked in the middle, while *µ*_*a*_ is ranked lower. On the optimal position they were ranked much higher. The median and maximum *µ*^∗^ on SoFPAN are respectively 0.14 and 1.71 mm^2^ (0.04 and 1.19 mm^2^) for 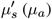. These parameters have some influence, but are subordinate to the other uncertain parameters such as subcellular location, expression level and cell morphology. Moreover, the need to include the light intensity profile in the neuron simulation was addressed by calculating the SoFPAN from excitation thresholds under uniform illumination, as well. Deviations of more than 20% are observed in more than 25% of the tested setups (soma excluded). Confinement to the basal dendrites result in the lowest percentage (18.17%) while the highest is achieved in case of the axon subcellular expression (34.60%). Overall, investigating optogenetic excitability under uniform field conditions provides a good initial approximation, but, accuracy drops for larger and asymmetrical section distributions. The latter was also observed in the strong pitch dependence of axon excitability in the pyramidal cells.

### D. Limitations and Future Work

This study focuses on single pulse excitation of CA1 cells. The occurrence of other spiking behavior is excluded. Likewise, when calculating the SoFPAN with high *I*_fiber_, a cell near the fiber end may exhibit bursting or depolarization block due to intense irradiance [40], [47]. Unlike with electrical stimulation, the photocurrent saturates for high light intensities (see figure 1 (C)). Therefore, extreme behavior is not expected when short pulses are applied. Additionally, the studied ChR2(H134R) opsin exhibits light-dark adaptation, i.e., the photocurrent is higher for a full dark adapted neuron but decreases towards a steady state value under prolonged illumination. The recovery time is in the order of seconds. During pulsed stimulation, the photocurrent of the first pulse will be higher than that of the subsequent pulses. Therefore, higher irradiances will be required to obtain reliable and pulse-locked spiking [45]. Furthermore, the capability of eliciting reliable spiking when opsin expression is confined to a specific subcellular region should be tested in future work. Additionally, this study focused on isolated cells. The interaction in neuronal networks will be of interest in future work, as well.

The effective level of opsin expression in-vivo is uncertain. Furthermore, to account for potential improvements in plasma membrane expression [3], [33], *G*_max_ is treated as a free parameter. However, due to the presence of inward rectification, it is unclear how the model parameter *g*_ChR2_ relates to the actual opsin expression level in-vivo. In our formulation of *G*(*V*), all proportionality factors are absorbed by *g*_ChR2_ (see equations (1) and (2)). While in the formulation of Grossman et al. (2011) and Williams et al. (2013), *G*(*V* = −68.83 mV) = 1 and *G*(*V* = −76.07 mV) = 1, respectively [36], [40]. These rectification functions cause a reduction of *g*_ChR2_ with a factor of 14.15 or 12.89 at those specific membrane potentials, compared to our formulation in equation (2).

As aforementioned, there is still some uncertainty on the tissue optical properties. Different studies have reported values that can differ up to an order of magnitude [25], [26], [63], [64]. Furthermore, brain tissue is binary classified as either gray or white matter, while tissue gradation is more continuous. The impact of the uncertainty of these parameters on the optogenetic excitability is tested in this study. However, it is investigated only locally near the parameters’ reported means (see table I), while the reduced scattering coefficient of white matter is reported to be 7-times higher than that of gray matter [26]. Additionally, tissue alterations due to foreign body reaction occur. A fibrous capsule is formed around the implanted fiber as reaction to blood-brain-barrier injury and gliosis caused by the presence of the implanted fiber itself [17]. In future work light propagation in a heterogeneous medium and its effect on excitability could be determined. Furthermore, exploring the propagation of light at different wavelengths, such as for exciting red-shifted opsins, would also be of interest.

New opsins, either natural or genetically engineered, are discovered on a yearly basis [23], [24], [61], [65], [66]. While this is generally beneficial, it can hinder the development of optogenetic tools. Dividing research among multiple opsins may limit the understanding of a single opsin’s interactions with neurons and its capabilities. The question arises whether the gathered insights here are transferable to other opsins as well. Previous studies have demonstrated the influence of channel kinetics on factors like irradiance thresholds, spike reliability, and behavior [23], [28], [40]. The exact values of *I*_th_ will thus differ for another opsin. However, these values are already uncertain due to multiple other uncertainties in other parameters like *G*_max_. Furthermore, these differences will affect all tested cases equally. Therefore, we expect that the observed trends and rankings regarding optogenetic excitability will be applicable across different opsins. In future work, a similar study could be performed focusing on optogenetic silencing with inhibitory opsins like GtACR2 and WiChr [66], [67].

Tissue illumination causes heating. To avoid permanent tissue damage, the local temperature increase cannot exceed 6 °C [18]. Moreover, behavioral changes are already possible at lower temperature changes (*>* 1 °C). Several neural parameters, e.g., capacitance, ion channel conductance, and transmitter release and uptake, have been shown to be temperature dependent [18], [21], [68], [69]. The reported SoFPAN values are for fiber intensities up to 1000 mW/mm^2^. After extrapolation of the change in temperature results reported in Acharaya et al. (2021; figure 3 (A)) [18], this intensity corresponds with an estimated temperature increase of 4.85 °C after 100 ms. Consequently, temperature-induced changes in optogenetic excitability should be included in future work.

## V. Conclusion

In this in-silico study, we examined the optogenetic excitability of four CA1 cells using ChR2(H134R). Our findings reveal that, for a fixed amount of opsin channels (*G*_max_), confining the opsin to specific neuronal membrane compartments significantly enhances excitability. This confinement leads to threshold reductions exceeding 64% and up to 100% gains in the surface of fiber positions for the activation of neurons. Additionally, we determined that the perpendicular orientation of the fiber relative to the somato-dendritic axis yields superior results. Furthermore, we observed substantial intercell variability, with differences in thresholds above 20%. The bistratified cell exhibited the least excitability, while pyramidal cell 1 demonstrated the highest excitability, especially when the opsin is confined to the basal dendrites. These findings highlight the importance of considering neuron degeneracy while developing optogenetic tools. By screening various uncertain parameters, we identified opsin location and *G*_max_ having the greatest impact on simulation outcomes. Our study showed the advantages of computational modeling coupled with light propagation. An increased excitability is seen with optimal fiber positioning, i.e., perpendicular to the somatic-dendritic axis and focus on the most excitable cell region. Spatial confinement and enhancements of opsin expression levels are promoted strategies to improve optogenetic excitability. However, it should be noted that uncertainty in these parameters limits determining the exact irradiance thresholds.

## Acknowledgment

R. Schoeters is a PhD Fellow of the FWO-V (FR) (Research Foundation Flanders, Belgium). T. Tarnaud is a postdoctoral fellow of the FWO-V. This work is supported by the BOF project SOFTRESET.

The computational resources (Stevin Supercomputer Infrastructure) and services used in this work were provided by the VSC (Flemish Supercomputer Center), funded by Ghent University, FWO and the Flemish Government – department EWI.

## Supplementary Info

**Fig. S1.**
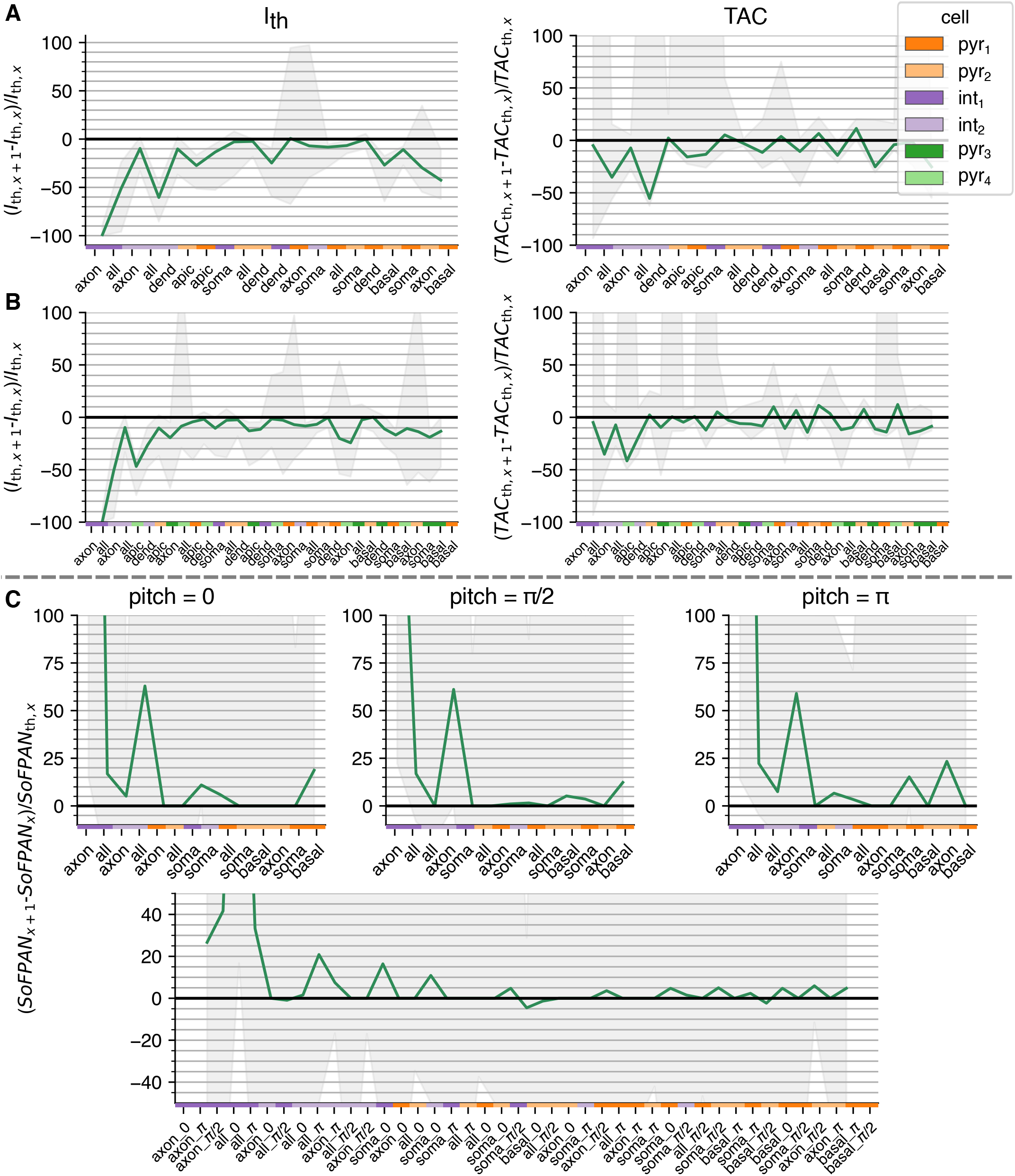
Median of relative change of ranked metrics. **A, B** Ranked *I*_th_ and TAC of the original four tested CA1 cells and with additional pyramidal cells, respectively, under a uniform intensity field. **C** Ranked SoFPAN: pitch specific at the top, all combined bottom. The gray shading indicates the total variation due to the pulse duration and opsin conductancy in all subplots

**Fig. S2.**
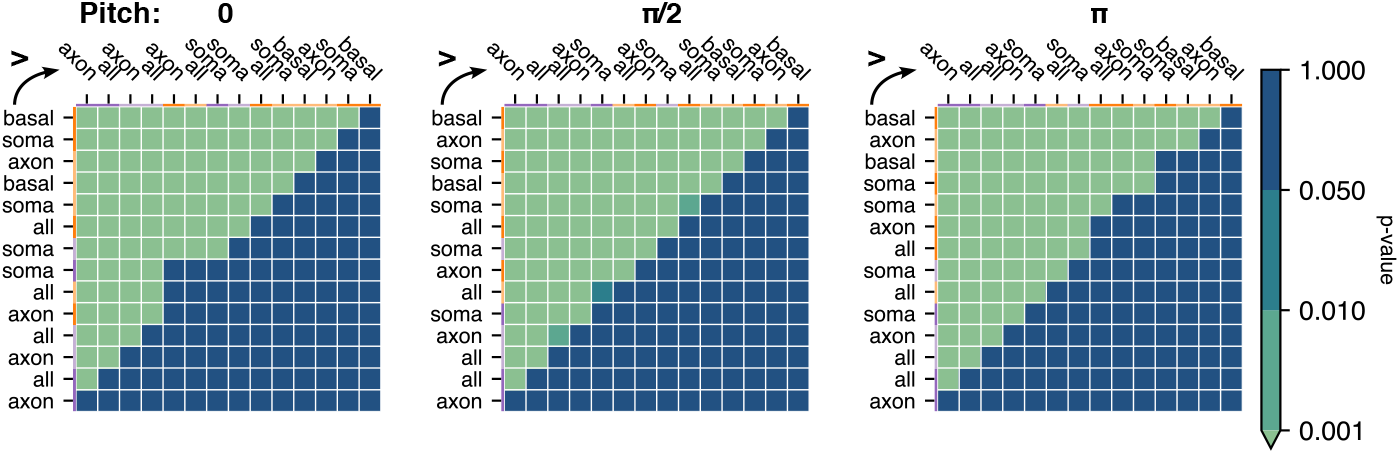
The p-value of the mutually, paired Wilcoxon signed-rank tests. Single-sided test if class at row is higher than class in the column for the SoFPAN for a given fiber pitch. The color code at the top and left indicate the cell model. Each population consists of 360 points (5 pd × 8 *G*_max_ 9 *I*_fiber_ values)

